# Light and dopamine impact two circadian neurons to promote morning wakefulness

**DOI:** 10.1101/2024.03.04.583333

**Authors:** Jasmine Quynh Le, Dingbang Ma, Xihuimin Dai, Michael Rosbash

## Abstract

In both mammals and flies, circadian brain neurons orchestrate physiological oscillations and behaviors like wake and sleep; these neurons can be subdivided by morphology and by gene expression patterns. Recent single-cell sequencing studies identified 17 *Drosophila* circadian neuron groups. One of these include only two lateral neurons (LNs), which are marked by the expression of the neuropeptide ion transport peptide (ITP). Although these two ITP^+^ LNs have long been grouped with five other circadian evening activity cells, inhibiting the two neurons alone strongly reduces morning activity; this indicates that they are prominent morning neurons. As dopamine signaling promotes activity in *Drosophila* like in mammals, we considered that dopamine might influence this morning activity function. Moreover, the ITP^+^ LNs express higher mRNA levels than other LNs of the type 1-like dopamine receptor Dop1R1. Consistent with the importance of Dop1R1, CRISPR/Cas9 mutagenesis of this receptor only in the two ITP^+^ LNs renders flies significantly less active in the morning, and *ex vivo* live imaging shows that dopamine increases cAMP levels in these two neurons; cell-specific mutagenesis of *Dop1R1* eliminates this cAMP response to dopamine. Notably, the response is more robust in the morning, reflecting higher morning Dop1R1 mRNA levels in the two neurons. As morning levels are not elevated in constant darkness, this suggests light-dependent upregulation of morning Dop1R1 transcript levels. Taken together with enhanced morning cAMP response to dopamine, the data indicate how light stimulates morning wakefulness in flies, which mimics the important effect of light on morning wakefulness in humans.

## Introduction

Wake and sleep are regulated by both a circadian and a homeostatic process.^1^ The latter monitors the duration an animal has been awake and the accumulation of sleep debt, whereas the 24-hour circadian drive determines when this sleep debt can be discharged. When circadian drive is high and sleep debt is low, animals transition more easily to wake. The detailed interactions between these two processes in driving circadian behavior remain unclear.

Circadian drive is usually reset each day by light. In mammals including humans, the light signal is passed from the retina to the suprachiasmatic nucleus (SCN) in the hypothalamus via the retinal-hypothalamic track. The SCN is considered the master clock and the most important circadian pacemaker in mammals.^2^ Additionally, light reduces sleep inertia, the phenomenon that describes grogginess upon transitioning to wake.^3^ Studies have also shown that premature light, illumination before scheduled waking, reduces sleep inertia.^4,5^ This is the case when we experience early morning sunlight during summertime or simulated dawn with artificial lights.

The circadian and homeostatic processes collaborate to craft the stereotyped bimodal activity pattern of *Drosophila melanogaster*, which peaks in the morning around the onset of dawn (or lights-on) and again in the evening around the onset of dusk (or lights-off). This activity pattern is dependent on an important subset of the 150 circadian neurons, the 15 pairs of lateral neurons (LNs) that are symmetrically present on both sides of the brain. They comprise of both morning cells and evening cells, which regulate locomotor activity in the morning and in the evening, respectively.^6,7^

The morning cells comprise of eight pairs of ventrally-located lateral neurons (LN_v_s), four large LN_v_s (lLN_v_s) and four small LN_v_s (sLN_v_s); the sLN_v_s also regulate rhythmicity in constant darkness.^8^ All eight LN_v_s express the neuropeptide pigment dispersing factor (PDF), which synchronizes the activity of many other circadian neurons.^9^ The seven pairs of evening cells consist of six dorsolateral neurons (LN_d_s) and one remaining sLN_v_; the latter cell is PDF^−^ negative and also known as the 5^th^ sLN_v_. This sLN_v_ and three of the six LN_d_s are sensitive to PDF from the morning cells. This is because these four cells express the PDF receptor (PDFR).^10^ They are also light sensitive as they express the blue light-sensitive circadian protein CRYPTOCHROME (CRY), which is required in the evening cells for light entrainment.^11^ The remaining three LN_d_s are both PDFR^−^ and CRY^−^.^12^

This substantial heterogeneity in evening cell gene expression is mirrored by a comparable heterogeneity in neuropeptide expression. Two of the CRY^+^ LN_d_s express both the short neuropeptide F (sNPF) and Trissin,^13^ whereas the third CRY^+^ LN_d_ expresses the neuropeptide NPF.^14,15^ Two CRY^−^ LN_d_s also express NPF as does the 5^th^ sLN ^14,15^ The 5^th^ sLN_v_ as well as one CRY^+^ LN_d_ also express the neuropeptide ion transport peptide (ITP).^16,17^ The ITP^+^ sLN_v_ and ITP^+^ LN_d_ (ITP^+^ LNs) contribute to the regulation of the evening locomotor activity peak.^15,17^

Recent electron microscopy (EM) connectome data highlights a comparable heterogeneity in evening cell connectivity.^18–21^ LN_d_s can be categorized into three groups based on shared connectivity patterns, two CRY^+^ groups and one CRY^−^. One of these CRY^+^ groups contains the two ITP^+^ LNs, which share strong synapses with each other,^19^ suggesting shared functions.

This dramatic heterogeneity parallels our previous characterization of circadian neuron transcriptomes at different clock times using bulk and single-cell RNA sequencing.^13,22,23^ The more recent single-cell RNA sequencing datasets used an unsupervised clustering method to identify high-confidence molecular subtypes with marker genes that match known identifiers of subtypes, such as G-protein coupled receptors and neuropeptides.^13,23^ This extensive dataset sits alongside years of immunohistochemistry work from various labs and underscores the molecular diversity of the evening neurons.

With a focus on understanding how these seven circadian neurons regulate sleep and wake, we investigated how they are affected by the arousal signal dopamine. Previous studies indicated that dopamine stimulates morning cell activity, specifically the lLN_v_s, which respond to dopamine through increases in cAMP signaling. Light suppresses this response by upregulating the inhibitory dopamine receptor Dop2R.^24^ Another study found that dopamine signaling to the lLN_v_s is mediated by the dopamine receptor Dop1R1and suggested that another dopamine receptor Dop1R2 present on the sLN_v_s promotes nighttime sleep.^25^ In yet another recent study, we showed that dopamine enhanced sleep through yet another circadian neuron group, the DN1s.^26^ However, a link between dopamine and the important wake-promoting evening cells has not been established. We were also encouraged to focus on the two ITP^+^ LNs in the context of sleep and wake because a genetic intersection strategy combined with morphological and molecular data shows that they are upstream of the other evening cells.

Indeed, optogenetic activation of novel and highly specific split-GAL4 lines indicates that the two ITP^+^ LNs can promote wakefulness by reducing sleep pressure and sleep depth. Relevant to dopamine, these cells express the wake-affiliated dopamine receptor Dop1R1 and CRISPR/Cas9-mediated mutation of this receptor only in the two ITP^+^ LNs significantly decreases activity in the morning. Since the Dop1R1 knockout primarily increases sleep depth, this suggests that dopamine signaling to these ITP^+^ LNs serves to decrease sleep depth and promote wakefulness in the morning. Consistent with this notion, *ex vivo* live imaging of cAMP in ITP^+^ LNs confirms that dopamine signaling to these neurons is mediated by Dop1R1. These results not only reveal an unexpected function of the ITP^+^ LNs, specifically their role in regulating wake and sleep in the morning, but also shed light on a new dopaminergic signaling pathway within the circadian network.

## Results

### ITP-expressing lateral neurons top the evening cell connectomic hierarchy

The evening cells are implicated in promoting the locomotor activity peak towards the end of the daytime and include six LN_d_s and one sLN_v_. Using the publicly available NeuPrint hemibrain and FlyWire whole brain electron microscopy connectome datasets,^18,20,21,27^ we compared connection strengths among all evening cells in two adult female *Drosophila* brains. The hemibrain connectome displays one set of evening cells (Figure 1A, left), while the FlyWire whole brain connectome shows two sets—one from each brain hemisphere (Figure 1A, right).

**Figure 1.**
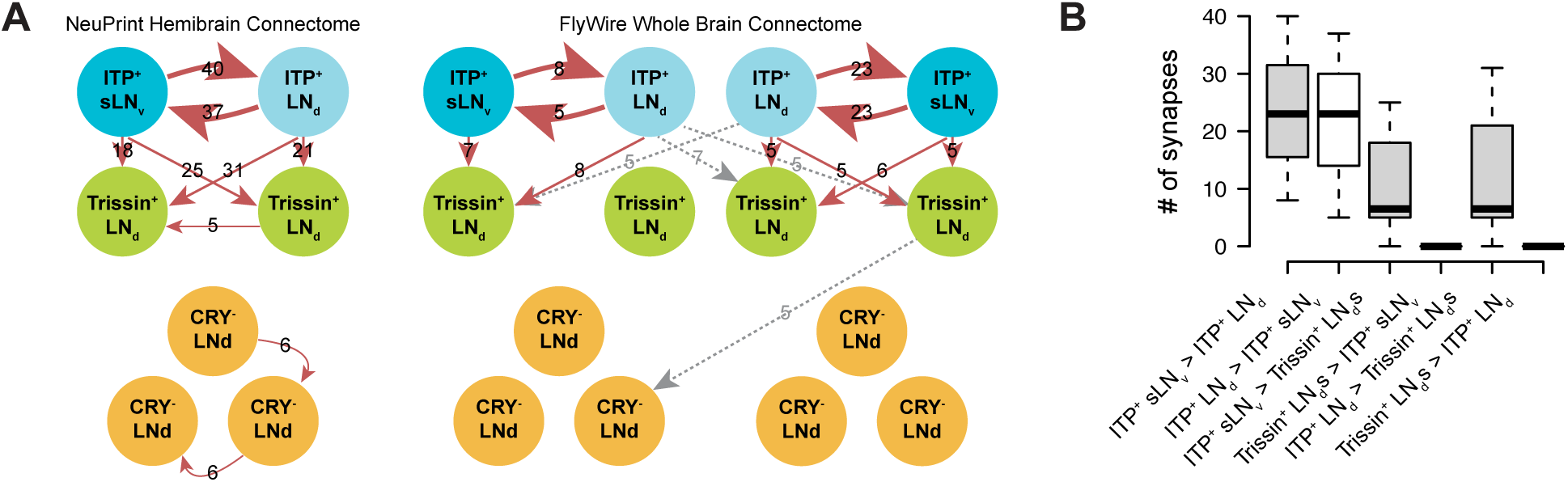
ITP-expressing lateral neurons top the evening cell connectomic hierarchy. (A) EM connectivity graphs showing connectivity weights of the evening cells with a threshold of at least 5 synapses of the NeuPrint Hemibrain Connectome^27^ (left) and the FlyWire Whole Brain Connectome^20,21^ (right). Direction of synapses are indicated by arrows and number of synapses are indicated by numbers. Gray dashed arrows indicate contralateral synapses. (B) Boxplot quantifying number of synapses between ITP^+^ sLN_v_s, ITP^+^ LN_d_, and Trissin^+^ LN_d_s in individual directions from both the NeuPrint Hemibrain Connectome^27^ and the FlyWire Whole Brain Connectome^20,21^ dataset.

Despite variations in the number of synapses between connectomes and even between hemispheres of the whole brain connectome, three consistent themes emerged across all three sets of evening cells. First, the ITP-expressing sLN_v_ and ITP-expressing LN_d_ (ITP^+^ LNs) exhibit strong interconnections, surpassing by far all connections with any other evening cells. Second, the ITP^+^ LNs synapse onto only two other evening cells, the two Trissin-expressing LN_d_s (Trissin^+^ LN_d_s). Notably, this connection is not reciprocated (Figure 1B), which reinforces this top-down view. Third, we did not observe any robust synaptic connectivity between the ITP^+^ LNs and the remaining three LN_d_s using a threshold of five synapses to define a valid connection. However, there are weak sub-threshold unidirectional connections from the ITP^+^ sLN_v_s and from one Trissin^+^ LN_d_ to the remaining CRY^−^ LN_d_s based on the NeuPrint hemibrain dataset.^19^ In summary, the synaptic weights and directions within all evening cells strongly suggest that the ITP^+^ LNs sit at the top of this circadian neuron group.

To provide finer detail in morphologically characterizing the ITP^+^ LNs, we employed a method that utilizes fluorescent labeling at the intersection of a GAL4 line and a LexA line (Figure S1A). By utilizing *Clk856-GAL4*^28^ to express a flippase-dependent fluorescent reporter *UAS-FRT-STOP-FRT-mVenus* in nearly all circadian neurons and an *ITP-LexA* knock-in line^29^ with *LexAop-flippase* to express flippase in all ITP-expressing neurons, we identified two neurons marked by both driver lines. Co-staining with an antibody for the circadian protein Period (PER) confirmed that a single LN_d_ and a single sLN_v_ are the only ITP^+^ circadian neurons (Figure S1B), consistent with previous ITP antibody staining findings.^17^ Notably, the ITP^+^ LN_d_ not only projects to the dorsal region of the brain but it also projects to the accessory medulla (AMe), unlike other evening cells. This is a key circadian region located at the anterior of the brain. It contains the LN_v_s and receives inputs from the eyes.^30–32^

Additionally, the axonal projection to the AMe from the ITP^+^ LN_d_ gives to a distinctive axonal bifurcation feature (Figure S1B, center, white arrowhead), which is absent in the other LN_d_s (Figure S1C, right). The bifurcation is also evident in EM reconstructions of the ITP^+^ LN_d_ (Figure S1C, center) but not in other evening cells (Figure S1C, right) or the ITP^+^ sLN_v_ (Figure S1C, left). The characteristic bifurcation allows for confident morphological distinction of the ITP^+^ LNd from other evening cells.

### ITP^+^ LNs are wake-promoting circadian neurons

Given the distinct molecular, morphological, and connectomic features of the ITP^+^ LNs, we asked whether these two neurons alone could influence circadian locomotor activity and sleep. To gain transgenic access to these neurons, we screened expression patterns of Janelia split-GAL4 driver lines from the Rubin Lab with the fluorescent protein expressed from *UAS-EGFP*. We looked for a driver line exhibiting the characteristic axonal bifurcation of the ITP^+^ LNd and identified a novel split-GAL4 driver line, *ss00639-GAL4*^33^ (Figure 2A): it only labels the two ITP^+^ LNs. There is also the *MB122B-GAL4*^34^ line, which labels the two ITP^+^ LNs and two other circadian neurons, the Trissin^+^ LNds (Figure S2D).

**Figure 2.**
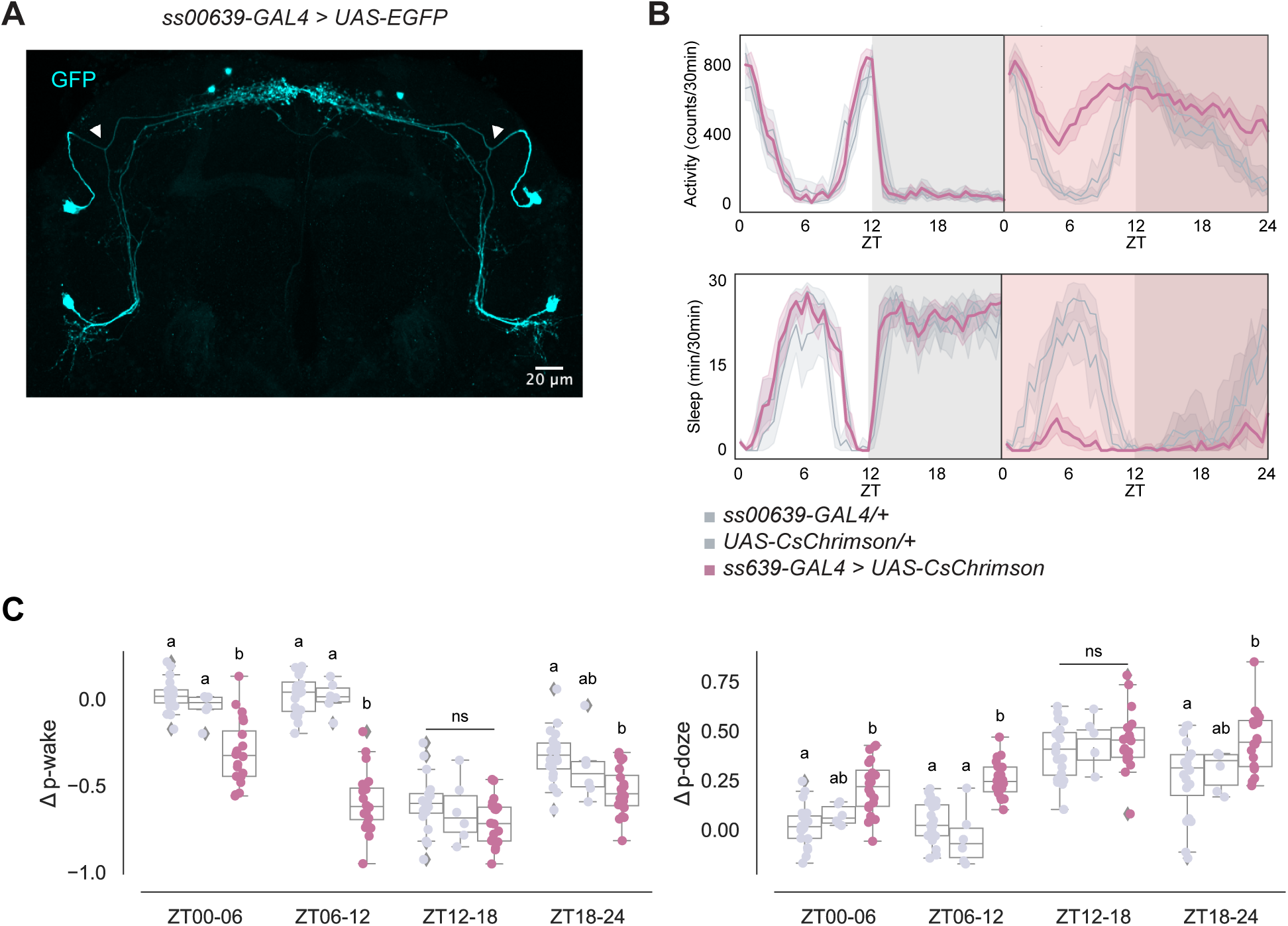
ITP^+^ LNs are wake promoting circadian neurons. (A) Representative whole-brain maximum intensity projection of *ss00639-GAL4* labeling ITP^+^ LNs with GFP (cyan) antibody staining. Axonal bifurcations of the ITP^+^ LN_d_s are marked by white arrowheads. (B) Timeseries plots of activity and sleep of female flies expressing red light-sensitive CsChrimson in ITP^+^ LNs at baseline (left) and with 24-hour red LED optogenetic activation (right) in 12:12 LD. Heterozygous *ss00639-GAL4* and *UAS-CsChrimson* controls are in gray. Experimental flies expressing both transgenes are colored. Bold lines are means and shaded regions are 95% confidence intervals of the means. (C) Boxplots quantifying the change in p-wake (left) and p-doze (right) of female flies expressing red light-sensitive CsChrimson in ITP^+^ LNs during the red LED optogenetic activation day from baseline day in six-hour time bins. Genotypes are depicted by the same color scheme as in (B). Letters represent statistically distinct groups as tested by a Kruskal Wallis test, post-hoc Mann Whitney U multiple comparisons method, and a Bonferroni-corrected significance value of p < 0.01667. Groups labeled with “ns” are not statistically distinct.

We then expressed a red light–sensitive channelrhodopsin, *UAS-CsChrimson*^35^, together with *ss00639-GAL4* to be able to optogenetically activate the ITP^+^ LNs with red light. The two parental lines with heterozygous expression of GAL4 and UAS alone were used as controls. A free-standing video-recording setup, FlyBox^34^ was used to observe locomotor and sleep behavior of these flies under standard 12:12 light-dark conditions (LD) and also delivered optogenetic stimulation.

During 12:12 LD baseline conditions, control flies and experimental flies expressing CsChrimson in the ITP^+^ LNs exhibited similar activity and sleep levels (Figure 2B, left panels). During 24 hours of red light stimulation however, experimental flies had dramatically higher activity levels and reduced sleep (Figure 2B, right panels) compared to control flies. Both control lines were unaffected by red light during the daytime, whereas optogenetically activated experimental flies lost most of their daytime sleep. Although control flies were woken by the stimulating red light during the nighttime, flies with both transgenes were much more active and had much lower nighttime sleep. This suggests that optogenetic activation of the ITP^+^ LNs promotes activity much more strongly than white light alone. Closer inspection of sleep structure during the daytime indicated ITP^+^ LNs activation increases the probability of sleeping flies waking up (p-wake) and decreases the probability of awake flies falling asleep (p-doze) during ZT00-06 and ZT06-12 (Figure 3C).^36^

**Figure 3.**
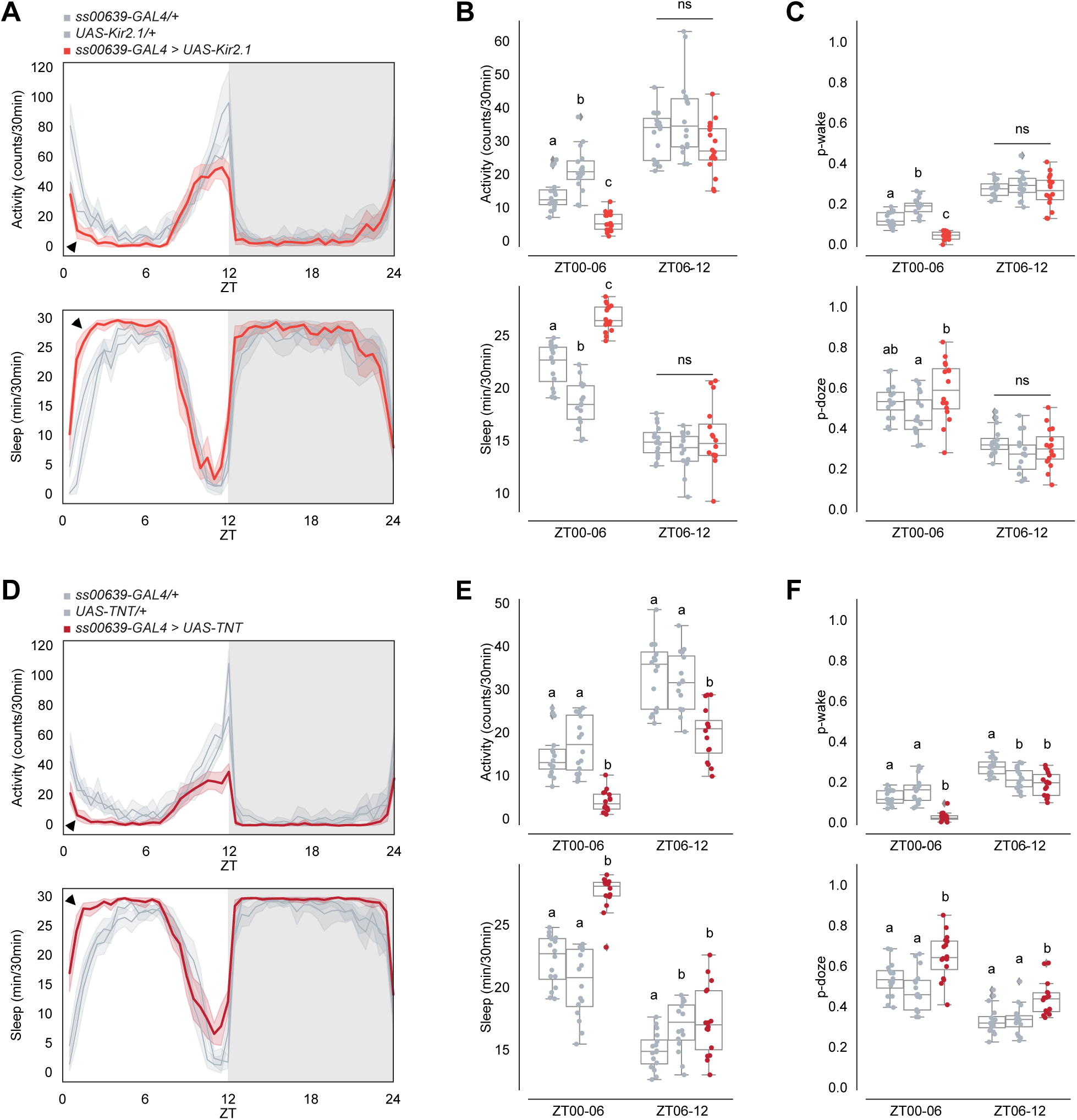
Inhibiting ITP^+^ LNs decreases early morning wakefulness. (A) Timeseries plots of activity (top) and sleep (bottom) of male flies expressing inhibitory Kir2.1 potassium channels in ITP^+^ LNs in 12:12 LD. Heterozygous *ss00639-GAL4* and *UAS-Kir2.1* controls are in gray. Experimental flies expressing all transgenes are colored. Bold lines are means and shaded regions are 95% confidence intervals of the means. Black arrowheads point to the major difference between control and experimental flies between ZT00 and ZT06. (B) Boxplots quantifying activity (top) and sleep (bottom) of male flies expressing inhibitory Kir2.1 potassium channels in ITP^+^ LNs during ZT00-06 and ZT06-12 daytime bins. Genotypes are depicted by the same color scheme as in (A). Letters represent statistically distinct groups as tested by a Kruskal Wallis test, post-hoc Mann Whitney U multiple comparisons method, and a Bonferroni-corrected significance value of p < 0.01667. Groups labeled with “ns” are not statistically distinct. (C) Boxplots quantifying p-wake (top) and p-doze (bottom) of male flies expressing inhibitory Kir2.1 potassium channels in ITP^+^ LNs during ZT00-06 and ZT06-12 daytime bins. Genotypes are depicted by the same color scheme as in (A). Letters represent statistically distinct groups as tested by a Kruskal Wallis test, post-hoc Mann Whitney U multiple comparisons method, and a Bonferroni-corrected significance value of p < 0.01667. Groups labeled with “ns” are not statistically distinct. (D) Timeseries plots of activity (top) and sleep (bottom) of male flies expressing synaptic transmission inhibitor TNT in ITP^+^ LNs in 12:12 LD. Heterozygous *ss00639-GAL4* and *UAS-TNT* controls are in gray. Experimental flies expressing all transgenes are colored. Bold lines are means and shaded regions are 95% confidence intervals of the means. Black arrowheads point to the major difference between control and experimental flies between ZT00 and ZT06. (E) Boxplots quantifying activity (top) and sleep (bottom) of male flies expressing synaptic transmission inhibitor TNT in ITP^+^ LNs during ZT00-06 and ZT06-12 daytime bins. Genotypes are depicted by the same color scheme as in (D). Letters represent statistically distinct groups as tested by a Kruskal Wallis test, post-hoc Mann Whitney U multiple comparisons method, and a Bonferroni-corrected significance value of p < 0.01667. Groups labeled with “ns” are not statistically distinct. (F) Boxplots quantifying p-wake (top) and p-doze (bottom) of male flies expressing synaptic transmission inhibitor TNT in ITP^+^ LNs during ZT00-06 and ZT06-12 daytime bins. Genotypes are depicted by the same color scheme as in (D). Letters represent statistically distinct groups as tested by a Kruskal Wallis test, post-hoc Mann Whitney U multiple comparisons method, and a Bonferroni-corrected significance value of p < 0.01667. Groups labeled with “ns” are not statistically distinct.

We next tested whether the Trissin^+^ LN_d_s are also activity-promoting like the ITP cells. Because we did not have a Trissin-only LN_d_ line, we activated these two neurons using an intersectional strategy. *Clk856-GAL4* labels all LNs and other circadian neurons, so we combined this GAL4 line with *UAS-FRT-STOP-FRT-CsChrimson*, *LexAop-flippase*, and a *Trissin-LexA* knock-in line^29^ to label only the two Trissin^+^ LN_d_s (Figure S2A). There was no change in activity, sleep, p-wake, nor p-doze observed when compared to heterozygous genetic controls (Figure S2B-C). However, activating both the ITP^+^ LNs and the Trissin^+^ LN_d_s with *MB122B-GAL4* (Figure S2D) resulted in a significant increase in activity and decrease in sleep (Figure S2E) not unlike the effect with *ss00639-GAL4* shown above. With *MB122B-GAL4*, p-wake and p-doze are affected only during ZT00-06 and not during ZT06-12 (Figure S2F). These data collectively indicate that the ITP^+^ LNs are wake-promoting but that the Trissin^+^ LN_d_s are not.

### Inhibiting ITP^+^ LNs decreases morning wakefulness

To complement the above activation experiments, we silenced the ITP^+^ LNs with two independent approaches: expressing the inward rectifying K^+^ channel Kir2.1 with *UAS-Kir2.1*^37^ and expressing tetanus toxin (TNT) with *UAS-TNT*^38^ to block neurotransmitter release with *ss00639-GAL4*.

Silencing with *UAS-Kir2.1* decreased activity and increased sleep during the daytime, particularly between ZT00 and ZT06, compared to heterozygous genetic controls (Figure 3A-B). P-wake is decreased between ZT00 and ZT06, whereas the change in p-doze is not significant in compared to both heterozygous genetic controls (Figure 3C). The results were very similar with *UAS-TNT* (Figure 3D-F). Both inhibition methods also caused a small decrease in the evening activity peak compared to controls, aligning with previous reports on evening cell inhibition^15,17,39^ although the effect is statistically significant with TNT, but not Kir2.1 (Figure 3B, E). Consistent results were also obtained observed with silencing and blocking neurotransmission of both ITP^+^ LNs and Trissin^+^ LN_d_s using *MB122B-GAL4* (Figure S3A-F). These findings further underscore the activity-promoting role of the two ITP^+^ LNs and focused our attention on the morning, particularly between ZT00 and ZT06, as well as the more expected effect on the evening activity peak.

### ITP^+^ LNs express higher levels of *Dop1R1* in the morning

Because the activation of different subsets of dopaminergic neurons has been demonstrated to promote wakefulness in *Drosophila*,^40^ we considered dopamine as a good candidate for an upstream arousal molecule. Moreover, our single-cell RNA sequencing data of circadian neurons^13^ showed that most circadian neurons express transcripts encoding a variety dopamine receptors (Figure 4A). For example, Dop1R1 and DopEcR are broadly expressed in circadian neurons, whereas Dop1R2 is limited to only a few clusters. The inhibitory receptor Dop2R is expressed at low levels in most clusters but more highly expressed in specific clusters. Expression within evening cells is visualized more easily by examining circadian time points in more detail. Dop1R1 and Dop2R show similar expression levels across all evening cells, while Dop1R2 is either absent or expressed at very low levels in all clusters (Figure S4A-C). DopEcR expression levels are high but variable in evening cells, consistent with other circadian neurons (Figure S4D).

**Figure 4.**
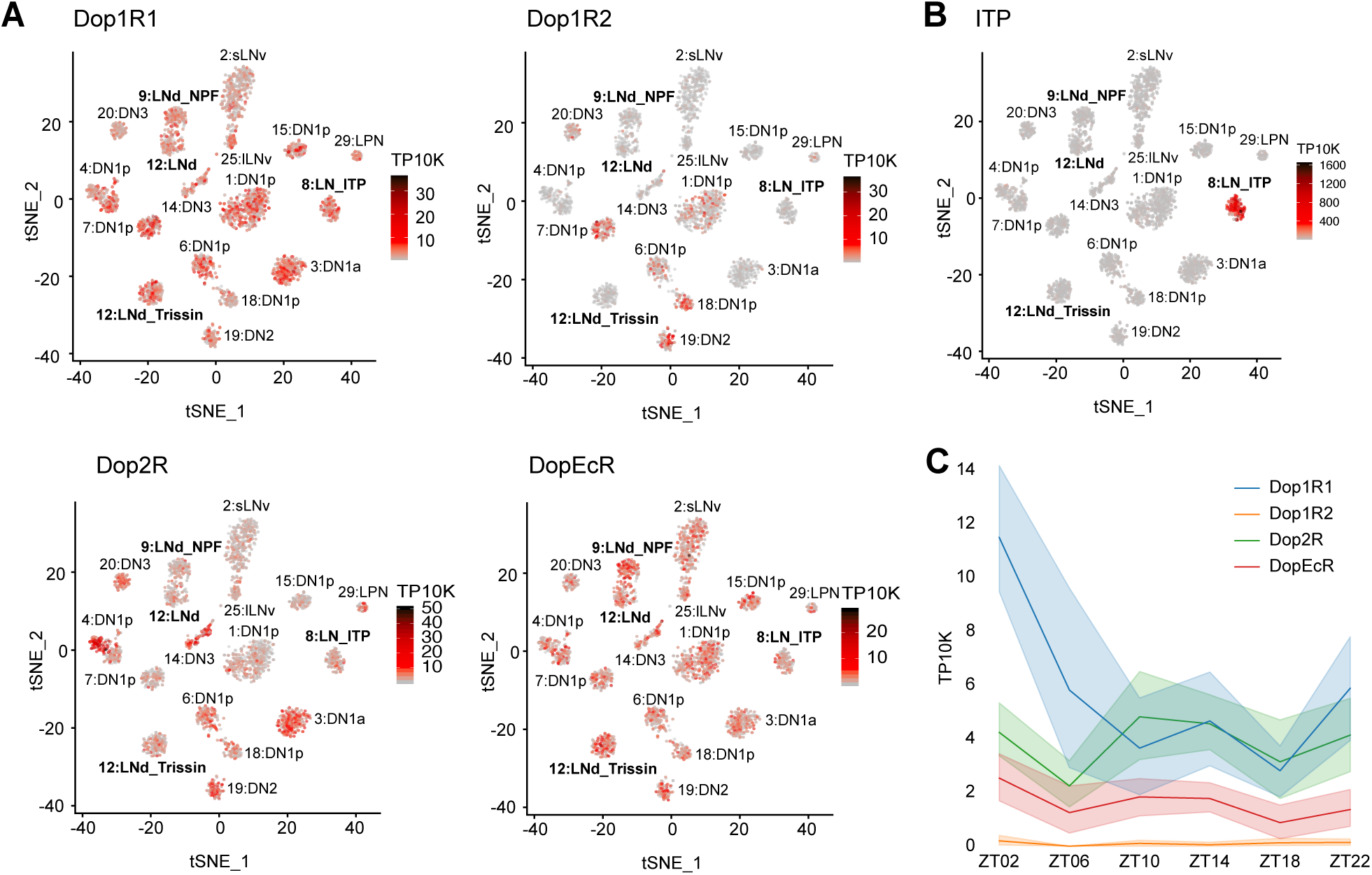
ITP^+^ LNs highly express *Dop1R1* mRNA. (A) Single-cell RNA sequencing data of circadian neurons^13^ displayed as tSNE plots and shaded by levels of dopamine receptor mRNA expression in number of transcripts per ten thousand transcripts. Each dot represents a single cell and each cluster represents a cell-type. Evening cell clusters marked by bold text. (B) Single-cell RNA sequencing data of circadian neurons^13^ displayed as a tSNE plot and shaded by levels of *ITP* mRNA expression in number of transcripts per ten thousand transcripts. Each dot represents a single cell and each cluster represents a cell-type. Evening cell clusters marked by bold text. (C) Timeseries plot of dopamine receptor mRNA expression levels around the clock in ITP^+^ LNs in number of transcripts per ten thousand transcripts. Different colors represent different dopamine receptors mRNAs. Bold lines are means and shaded regions are 95% confidence intervals of the means.

The ITP^+^ LNs are recognizable through their notably high ITP expression (Figure 4B). These two cells express Dop1R1 prominently, along with some Dop2R and DopEcR, while virtually lacking Dop1R2 expression (Figure 4A). Intriguingly, Dop1R1 mRNA in the ITP^+^ LNs follows a cycling pattern with a peak in the morning at ZT02 (Figure 4C). Intriguingly, this is the only evening cell cluster and dopamine receptor for which morning expression is deceased in constant darkness, indicating a special relationship between light, the ITP^+^ LNs and Dop1R1 (Figure S4A).

### Functional knock-out of Dop1R1 in ITP^+^ LNs decreases morning wakefulness

Because of the morning sleep/activity effect of activation and inhibition shown above, we further pursued the contribution of dopamine and Dop1R1 to the two ITP^+^ LNs. To this end, we employed a CRISPR/Cas9 mutagenesis method^26,33,41^ to specifically knock out Dop1R1 function in wake-promoting ITP^+^ LNs. Given that Dop1R1 is known as an excitatory dopamine receptor,^42^ our hypothesis was that knocking out Dop1R1 in these neurons would diminish their activity, leading to reduced wake-promotion throughout the day. However, the results revealed a decrease in activity and an increase in sleep only in the morning, specifically between ZT00 and ZT06 under standard 12:12 LD conditions (Figure 5A-B). Importantly, activity and sleep levels remained indistinguishable from the control strains thereafter. Similar effects were observed with RNAi knock-down of Dop1R1 in these neurons (Figure S5C, black arrowheads). Given that ITP^+^ LNs are a subset of the evening cells responsible for regulating the evening activity peak,^15,17,39^ the unchanged evening activity anticipation and peak with Dop1R1 knock-out in these cells indicates a dedicated contribution of these cells, their Dop1R1 expression and dopamine to morning wakefulness.

**Figure 5.**
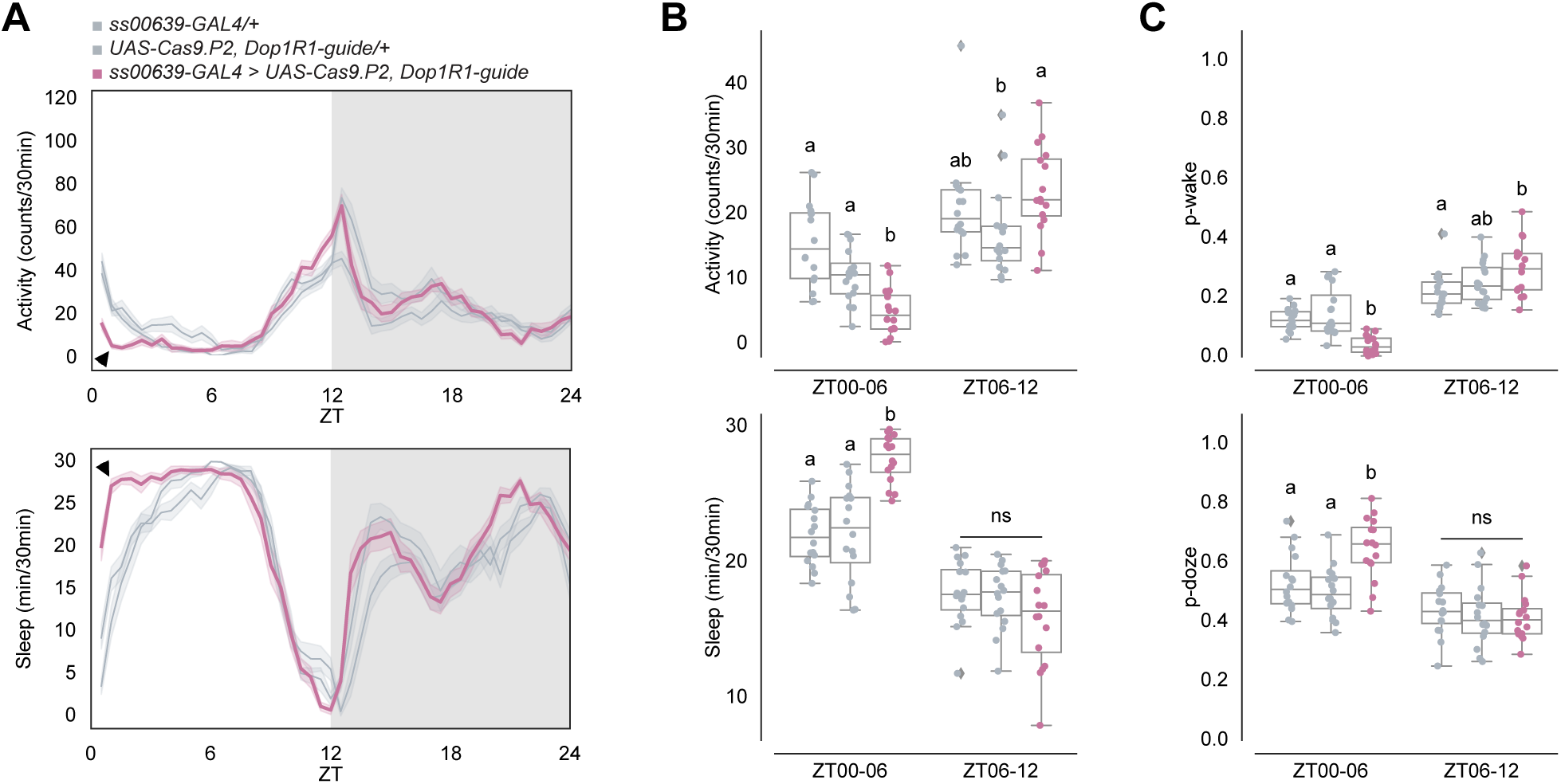
Functional knock-out of *Dop1R1* in ITP^+^ LNs decreases early morning wakefulness and increases sleep. (A) Timeseries plots of activity (top) and sleep (bottom) of male flies with *Dop1R1* knock-out in ITP^+^ LNs in 12:12 LD. Heterozygous *ss00639-GAL4* and *UAS-Cas9.P2, Dop1R1-guide* controls are in gray. Experimental flies expressing all transgenes are colored. Bold lines are means and shaded regions are 95% confidence intervals of the means. Black arrowheads point to the major difference between control and experimental flies between ZT00 and ZT06. (B) Boxplots quantifying activity (top) and sleep (bottom) of male flies with *Dop1R1* knock-out in ITP^+^ LNs during ZT00-06 and ZT06-12 daytime bins. Genotypes are depicted by the same color scheme as in (A). Letters represent statistically distinct groups as tested by a Kruskal Wallis test, post-hoc Mann Whitney U multiple comparisons method, and a Bonferroni-corrected significance value of p < 0.01667. Groups labeled with “ns” are not statistically distinct. (C) Boxplots quantifying p-wake (top) and p-doze (bottom) of male flies with *Dop1R1* knock-out in ITP^+^ LNs during ZT00-06 and ZT06-12 daytime bins. Genotypes are depicted by the same color scheme as in (A). Letters represent statistically distinct groups as tested by a Kruskal Wallis test, post-hoc Mann Whitney U multiple comparisons method, and a Bonferroni-corrected significance value of p < 0.01667. Groups labeled with “ns” are not statistically distinct.

The decrease in morning wakefulness can result from either a stronger drive towards sleep and/or a decreased wake drive. Examining sleep structure indicated that the decrease is due to both, a significant decrease in p-wake and an increase in p-doze between ZT00 and ZT06 compared to the control strains (Figure 5C). In contrast, the difference in the second half of the daytime, during ZT06 and ZT12 was more variable and not significant. Although activity and sleep were similarly affected with CRISPR/Cas9 knock-out (Figure S5A-B) and RNAi knock-down (Figure S5D, black arrowheads) in ITP^+^ LNs and Trissin^+^ LN_d_s with *MB122B-GAL4*, p-wake was significantly decreased between ZT00 to ZT06 while p-doze did not significantly differ from both controls (Figure S5C). These findings suggest that dopamine inputs to Dop1R1 in ITP^+^ LNs typically create a strong drive towards wakefulness by decreasing sleep depth in the morning.

### Dopamine increases cAMP levels in ITP^+^ LNs

Given the significance of Dop1R1 function in ITP^+^ LNs for morning wakefulness, we examined the cellular effects of dopamine on these neurons using an intracellular cAMP sensor, EPAC-H187^43^. We cloned this sensor into flies under UAS control^44^ and expressed it in ITP^+^ LNs using *ss00639-GAL4*. Live cAMP changes in response to dopamine were recorded in whole brain explants using confocal microscopy. To minimize network effects, brains were incubated in adult hemolymph-like saline (AHL) with tetrodotoxin (TTX) for at least five minutes before recording to inhibit polysynaptic inputs; all experiments were conducted with TTX. One minute of baseline activity was recorded before perfusing each brain with dopamine and with forskolin directly afterwards as a positive control.

In both the ITP^+^ LN_d_ and the ITP^+^ sLN_v_, we observed increases in cAMP in response to 400 uM dopamine and 50 uM forskolin (Figure S6A-B, blue traces). To confirm that the response to dopamine is mediated by Dop1R1, we co-expressed our sensor *UAS-EPAC-H187* and the Dop1R1-guides (with *UAS-Cas9.P2*) in *ss00639-GAL4* to measure the cAMP response to dopamine without functional Dop1R1 receptors. There were no responses to dopamine in either the ITP^+^ sLN_v_ or the ITP^+^ LN_d_ with the knock-out (Figure S6A-B, orange traces), indicating that the cAMP response is indeed driven through the Dop1R1 receptors. Importantly, both cells still responded to forskolin, showing that they can still mount a cAMP response through other pathways.

We hypothesized that cAMP responses to dopamine would be more pronounced during the light phase around ZT02 when *Dop1R1* mRNA levels are highest compared to the dark phase when *Dop1R1* mRNA levels are lowest (Figure 4C). To capture time-of-day differences in cAMP responses, we utilized unpublished 10XUAS versions of newly available and more sensitive EPAC sensors called cAMPFIRE.^45^

Robust responses to 100 uM dopamine and 50 uM forskolin were observed during the light phase between ZT01 and ZT03 and the dark phase between ZT13 and ZT15 in both ITP^+^ LNs with these *10XUAS-cAMPFIRE* sensors (Figure 6A-B). However, the magnitude of the response was significantly greater in the light phase than in the dark phase only in the ITP^+^ sLN_v_ (Figure 6C). The response in the ITP^+^ LN_d_ also trended higher during the light phase, but this difference was not statistically significant. The data indicate that the single ITP^+^ sLN_v_ may play a more substantial role than the ITP^+^ LN_d_ in dopaminergic modulation of morning wakefulness and suggest that the cycling of *Dop1R1* mRNA levels within these cells contributes to the control of time-of-day wakefulness.

**Figure 6.**
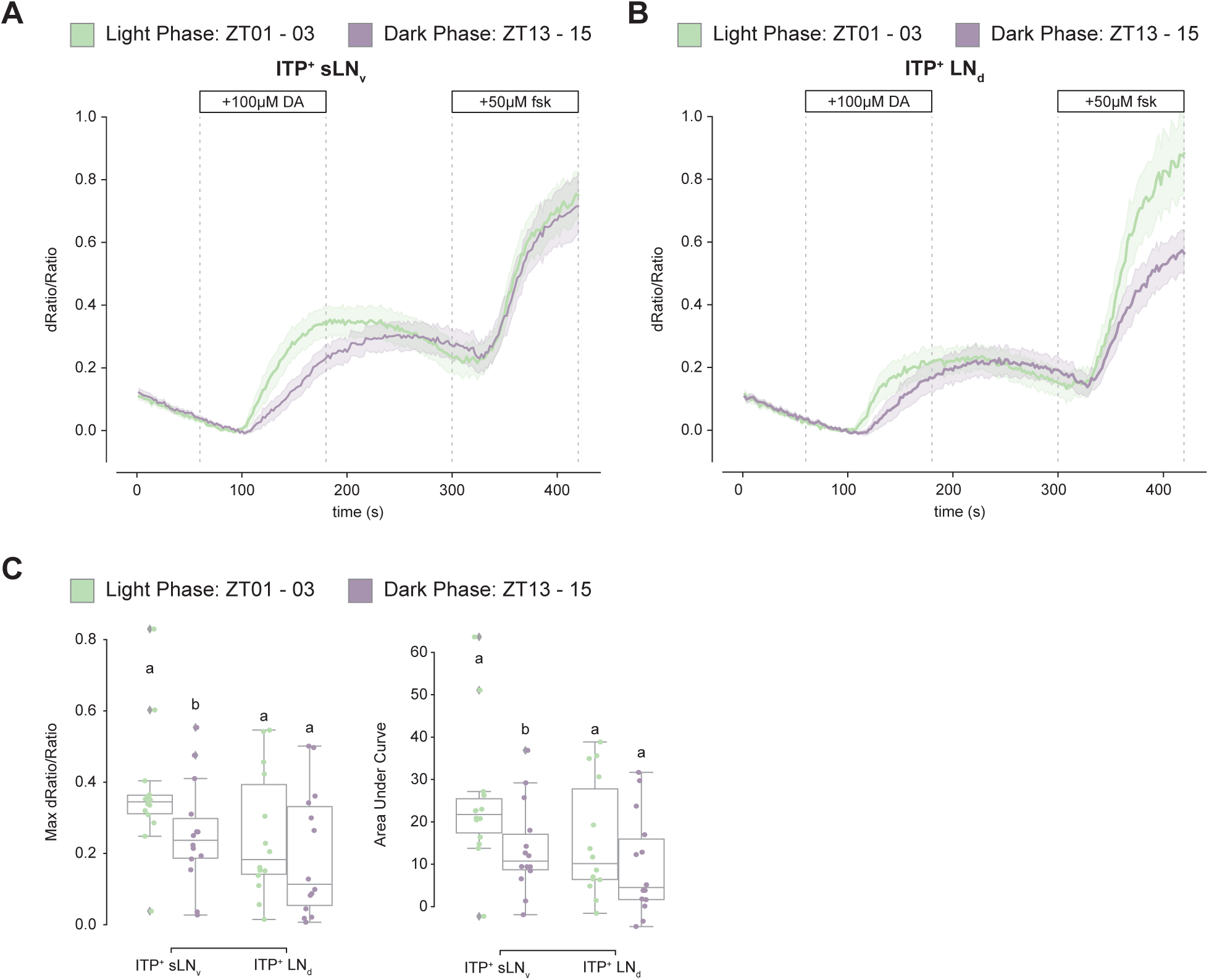
cAMP responses to dopamine are more robust during the day than the night. (A) Timeseries plot of cAMP levels at baseline and in response to 100µM dopamine and 50µM forskolin perfusion as measured by *10XUAS-cAMPFIRE-M* in the ITP^+^ sLN_v_ during the light phase (green) and dark phase(purple). Bold lines are means shown with a simple moving average of two data points and shaded regions are 68% confidence intervals of the means. (B) Timeseries plot of cAMP levels at baseline and in response to 100µM dopamine and 50µM forskolin perfusion as measured by *10XUAS-cAMPFIRE-M* in the ITP^+^ LN_d_ during the light phase (green) and dark phase(purple). Bold lines are means shown with a simple moving average of two data points and shaded regions are 68% confidence intervals of the means. (C) Boxplots quantifying the maximum dRatio/Ratio (left) and area under curve (right) of the ITP^+^ sLN_v_ and ITP^+^ LN_d_ in response to 100µM dopamine perfusion. Letters represent statistically distinct groups with p-value < 0.05 as tested by a Mann Whitney U test of individual cell-types.

## Discussion

Although the circadian evening neurons have been shown to be activity-promoting,^6,34^ there is little information that distinguishes the function(s) of one cell from another. To this end, we began by identifying and comparing the connectivity patterns of these seven neurons. The two ITP^+^ LNs emerged at the top of a hierarchy, hinting that they may have a special and important role. Subsequent identification of split-GAL4 lines for labeling ITP^+^ LNs showed that they promote wake at all times of day, but inhibition of these neurons led to a significant reduction only in morning wakefulness. Single-cell RNA sequencing showed that Dop1R1 receptor mRNA is more highly expressed in the ITP^+^ LNs than in other LNs. Dopamine also induces increased cAMP levels within these neurons, which is more pronounced during the day than the night. Furthermore, *Dop1R1* mRNA levels in ITP^+^ LNs are highest in the morning during light-dark conditions but not constant darkness, which suggests that light-dependent cycling of Dop1R1 is contributes to their morning arousal function.

Although the ITP^+^ LNs emerged as prominently interconnected as well as connected to other evening cells, this relationship was unidirectional. There were almost no pre-synaptic connections from the other evening cells to the ITP^+^ LNs. This conclusion was drawn after analyzing two EM connectome datasets and also builds upon earlier studies that primarily addressed inter-LN_d_ connections.^19^ In addition, the morphological distinctions between the four LN_d_ subtypes we identified was consistent with extensive prior research that classified LN_d_s based on their protein and RNA expression.^12,13,16^

The two ITP^+^ LNs have robust synaptic inputs into the two Trissin^+^ LN_d_s (see Figure 1A), which are a second evening cell subtype and are located directly downstream of the ITP^+^ LNs. These Trissin^+^ LN_d_s are included in the expression pattern of *MB122B-GAL4* along with the two ITP^+^ LNs. Because manipulating the activity of this driver had very similar effects to manipulating the activity of the two ITP^+^ LNs alone with *ss00639-GAL4*, the data suggest that there is little or no effect of activating only the Trissin^+^ LN_d_s. Indeed, directly activating only the Trissin^+^ LN_d_s, (Figure S2C), confirmed that the wake-promoting effect of *MB122B-GAL4* is due to the ITP^+^ LNs and not the Trissin^+^ LN_d_s.

Although the ITP^+^ LNs are clearly wake-promoting when activated, the effect of inhibition with two different methods was much more discrete. This suggests that activation likely induces “ectopic” effects, by swamping normal activity levels as well as normal time of day regulation. This is most easily interpreted by proposing that the ITP^+^ LNs can activate other evening cells, which are primarily responsible for evening locomotor activity. This is consistent with the connectomic data (Figure 1). However, inhibition of the ITP^+^ LNs did cause a small decrease in the evening activity peak, although it was not statistically significant with Kir2.1 inhibition (Figure 3B). This suggests that the ITP^+^ LNs do influence evening activity but that they are not the major neurons for this function.

The modest and discrete effect of ITP^+^ LN inhibition on morning locomotor activity reminded us of the temporal dynamics of *Dop1R1* transcript expression, which has a morning peak in the ITP^+^ LNs (Figure 4C). Because *Dop1R1* is also more expressed in ITP^+^ LNs compared to other evening cells, we considered dopamine as a good candidate for a relevant upstream molecule (also refer to Figure S4A). To address this possibility, we used CRISPR/Cas9 to knock out *Dop1R1*. This caused a substantial loss of wakefulness and increased sleep during the morning hours, notably between ZT00 and ZT06 (Figure 5A), which resembled the effect of neuronal inhibition. This sleep increase was predominantly due to a reduced likelihood of spontaneously awakening from a sleep state (Figure 5B), indicating that Dop1R1 plays a prominent role in enhancing morning arousal. This finding is concordant with prior studies demonstrating the wake-promoting effects of dopaminergic neurons.^40,46^

However, the overall impact of dopamine on the *Drosophila* circadian network is more complicated than its focused effect described here. Although dopamine also enhances locomotor activity through its effect on lLN_v_s,^24^ it has been reported to promote rather than inhibit sleep through its action on sLN_v_s and DN1s.^25,26^ Despite these other results, it is hard to ignore the similar effects of neuronal inhibition and the Dop1R1 knockout; they both indicate that dopamine impacts the ITP^+^ LNs to promote morning arousal.

This period of time, between ZT00-ZT06, is not normally associated with evening cell activity. This recalls the fact that the ITP^+^ LN_d_ cell is the only evening neuron to project to the AMe, where the PDF^+^ LN_v_s morning cells reside. The other ITP^+^ LN is the single PDF^−^ sLN_v_. Its cell body resides in the AMe adjacent to the PDF^+^ LN_v_s, i.e., both ITP^+^ LNs are well-positioned to receive information from, and/or send information to, the morning cells. Moreover, the ITP^+^ LNs express PDFR,^12^ making it very likely that these neurons are impacted by PDF expressed from the AMe-resident morning cells. Considering the known activity peaks of morning cells (sLN_v_s, late evening, ~ZT23; lLN_v_s, morning, ~ZT06), dopamine might effectively amplify PDF signaling to the two ITP^+^ LNs and thereby enhance the morning amplitude of circadian locomotor activity. All of these considerations indicate that these two evening cells are really morning cells and suggests that the evening cells should really be renamed activity cells, morning activity for the ITP^+^ LNs and evening activity for other LNs, most likely the three CRY-LNds. This suggestion is based on the strong effect on evening activity from the *dVPDF-GAL4* driver, which expresses in the two ITP^+^ LNs as well as the three CRY^−^ LN_d_s.^39^ Unfortunately, this remains speculation as there is no driver that only expresses in the three CRY^−^ LN_d_s.

The substantial cycling of *Dop1R1* mRNA levels in light-dark (LD) conditions is largely absent in constant darkness (DD). Intriguingly, the ITP^+^ LNs are the only evening cell cluster and *Dop1R1* mRNA the only dopamine receptor transcript for which morning expression is decreased in constant darkness, suggesting a special relationship between light and the Dop1R1/ITP^+^ LNs (Figure S4A).

A mechanistic explanation for the light-mediated upregulation of these mRNA levels in only these two neurons is of interest but unknown. However, a likely teleological explanation for this relationship is that light and dopamine amplify circadian morning wake-drive. This emphasizes the extent to which these two circadian neurons and their discrete behavioral output are sensitive to environmental conditions (light) as well as possibly internal state (dopaminergic tone). This relationship may also serve as a possible model for humans experiencing less sleep inertia, more wake-drive, with light exposure, either from early morning sun or from simulated dawn with artificial light before an alarm.^4,5^

Although the cycling of *Dop1R1* mRNA levels is based only on RNA sequencing data, the bigger dopamine-mediated cAMP response from daytime than from nighttime brains suggests that *Dop1R1* protein levels and activity in the two ITP^+^ LNs are also higher in the daytime than the nighttime. This is the first indication that the cycling of GPCR mRNA levels^13,22,26^ is of functional significance. Notably, the cycling of *Dop1R1* RNA levels and protein/activity levels are likely not identical in the two neurons, a possibility that is supported by the larger sLN_v_ day-night difference than LN_d_ day-night difference in response to dopamine perfusion. This suggests higher *Dop1R1* activity and perhaps a bigger morning arousal role for the ITP^+^ sLN_v_ than the ITP^+^ LN_d._ This distinction also serves as a reminder that even a two cell-cluster may harbor internal heterogeneity. More generally put, even the considerable cell type complexity of the fly brain circadian neuron system^13,19^ is likely to be an underestimate: every circadian neuron may be discrete, at the transcriptomic and anatomical level and perhaps even functionally.

## Acknowledgements

Many thanks to members of the Rosbash Lab for helpful discussions and help with this work, especially, Dr. Shlesha Richhariya and Dr. Kate Abruzzi. We thank our former colleague Dr. Fang Guo for establishing optogenetic experiments in the lab. Thank you very much also to Mohamed Adel from the Griffith Lab at Brandeis University particularly for help with imaging analysis and former member Dr. Timothy Wiggin for a lot of initial help with behavior analysis. Thank you to Dr. Heather Dionne, Dr. Aljoscha Nern, Dr. Gerry Rubin from Janelia Research Campus for generously sharing unpublished fly lines. We also thank the Yi Rao lab for sharing Chemoconnectome lines with us and the Haining Zhong lab for sharing unpublished cAMPFIRE constructs with us. This work was supported by the Howard Hughes Medical Institute, the National Institute of Mental Health Neuroscience Training Grant (NIH grant no. MH019929), and the National Institute of General Medical Sciences Genetics Training Grant (NIH grant no. GM007122).

## Materials and Methods

### Fly strains and husbandry

*Drosophila melanogaster* strains were reared on a standard cornmeal/agar medium supplemented with yeast under 12:12LD cycles at 25°C. Heterozygous genetic controls used in behavioral experiments were parental strains crossed to w1118 wild-type flies. Females used in all experiments were mated. Strains used in this study are listed in Table S1.

### Generation of fly line

To generate UAS-EPAC-H187 flies, we used NotI and XbaI to digest EPAC-S-H187 (Addgene #170348)^43^ and ligated it to a NotI- and XbaI-digested vector of pJFRC7-20xUAS (Addgene #26220)^44^. This was injected into attP1 on the 2^nd^ chromosome and vk00027 on the 3^rd^ chromosome (Rainbow Transgenic Flies, Inc.).

### Immunohistochemistry

Whole flies were fixed in PBS with 4% paraformaldehyde and 0.5% Triton X-100 for 2.5 hours while rotating at room temperature. The flies were then washed in PBS with 0.5% Triton X-100 (PBST) for 3x 10 minutes before their brains were removed via dissection. Dissected brains were washed 3x 15 minutes and blocked with 10% normal goat serum (NGS; Jackson Labs) in 0.5% PBST (blocking buffer) for 2 hours at room temperature or overnight at 4°C. Brains were then incubated at 4°C overnight in blocking buffer with the following antibodies and concentrations: chicken anti-GFP (1:1000; Abcam ab13970), rabbit anti-per (1:1000; Rosbash Lab), and rabbit anti-dsRed (Takara Bio 632393; 1:200). Then, brains were incubated with secondary antibodies (Alexa Fluor 488-conjugated anti-chicken, AlexA Fluor 633-conjugated anti-rabbit) diluted at 1:500 in blocking buffer for two hours at room temperature, and washed 3x 15 minutes with 0.5% PBST. Stained brains were mounted in VectaShield mounting medium (Vector Laboratories, Newark, CA). Images were acquired using a Leica Stellaris 8 confocal microscope or a Leica SP5 confocal microscope equipped with a white-light laser and a 40X oil objective, and processing was done using Fiji ^47^.

### EM Connectome Analysis

Hemibrain EM connectome data ^18^ was accessed and analyzed with NeuPrint.^27^ To identify our neurons of interest, we used the “Find Neurons” function and recorded their BodyIDs. To compare morphologies, we entered BodyIDs into the “Skeleton” visualization function. To look at connectivity strengths, we entered BodyIDs into the “Connectivity Graph” function.

Whole brain EM connectome data ^21^ was accessed and analyzed with FlyWire.^20^ To identify our neurons of interest, we used the “Search Cells and Annotations” function and recorded their IDs. To look at connectivity, we entered IDs into the “Network” function.

### Optogenetic Locomotor Behavior Assay

Optogenetic experiments were done in FlyBoxes, as previously described.^34^ Flies were aged 2-5 days old and loaded into individual wells of white 96-well plates containing 300uL of food with 5% sucrose, 2% agar, and 400uM all trans-retinal (Sigma Aldrich, R2500). Within a FlyBox, the plate is illuminated from underneath with infrared light supplied by an 850 nm LED board (LUXEON). Images of loaded plates were captured every ten seconds using a down-facing USB webcam (Logistic C910 with infrared light filter removed) placed at the top of the box and with Image Capture software. Entrainment light with a 12:12 LD cycle was provided by a white LED strip set to the minimum brightness. Optogenetic stimulation was given using high power red (630 nm) LEDs pulsing at 5Hz (0.08mW/mm^2^). All lights were controlled with an Arduino in the FlyBox. Each experiment included two full entrainment days and one baseline day before optogenetic manipulation. All experiments were conducted at ambient room temperature. Fly locomotor activity was extracted from the images using pySolo^48^ and pre-processed with DAMFileScan (Trikinetics, Waltham, MA).

### Standard Locomotor Behavior Assay

Standard experiments were done using the Drosophila Activity Monitor (DAM) system (Trikinetics, Waltham, MA), as previously described.^49^ Flies were aged 2-5 days old and loaded into individual glass tubes with 5% sucrose and 2% agar food on one end and a stopper on the other. Glass tubes with flies were then loaded onto DAMs, which recorded the number of beam crosses a fly makes. All experiments were conducted with 12:12 LD in 25°C incubators. Flies were entrained for two full days, and activity and sleep were averaged across the 2-3 days thereafter. Data was pre-processed with DAMFileScan (Trikinetics, Waltham, MA).

### Behavior Analysis

Activity and sleep data from both optogenetic and standard behavior experiments were analyzed with the Sleep and Circadian Analysis MATLAB Program (SCAMP, Vecsey Lab). P-wake and p-doze analyses were done using the Fly Sleep Probability analysis package.^36^ Error bars and shaded regions in the time series plots are 95% confidence intervals of the means. For all box plots, center lines indicate medians, box limits indicate 25^th^ and 75^th^ percentiles, and whiskers extend 1.5 times the interquartile range. Statistical significance of box plots were tested using one-way ANOVA with Tukey’s multiple comparisons test. All plots were created with Python, MATLAB, or R.

### Single-Cell RNA Sequencing

Single-cell RNA sequencing experiments and analysis were done by and reported in Ma et al., 2021. Briefly, flies expressing a fluorescent protein under the control of *Clk856-GAL4* were entrained and dissected at timepoints every four hours around the clock. Brains were dissociated into a single-cell suspension and sorted using a fluorescence-activated cell sorting (FACS) machine (BD Biosciences, Franklin Lakes, New Jersey). Single-cell RNA library preparation was done using a modified version of CEL-Seq2.^50,51^ The resulting data were aligned to Drosophila genome 6 (dm6) using zUMIs^52^ and STAR.^53^ Dimensional reduction, clustering, and differential gene expression analysis were all done using Seurat.^54^

### *Ex Vivo* EPAC Functional Imaging

UAS-EPAC-H187 or 10XUAS-cAMPFIRE-M were expressed in ITP^+^ LNs using *ss00639-GAL4*. Flies aged 0-5 days old were collected and entrained in 12:12 LD for at least two full days. Flies were collected in the light phase between ZT01 and ZT03 or in the dark phase between ZT13 and ZT15 for testing. Adult female brains were dissected in adult hemolymph-like saline (AHL; 108 mM NaCl, 5mM KCl, 2 mM CaCl_2_, 8.2 mM MgCl_2_, 4 mM NaHCO_3_, 1 mM NaH_2_PO_4_, 5 mM trehalose, 10 mM sucrose, 5 mM HEPES; pH 7.5) (Cold Spring Harbor Protocols, 2013) and 1 uM tetrodotoxin (AHL-TTX). Brains were mounted onto a poly-l-lysine-coated cover slip (Neuvitro Corporation, Camas, WA) on a SYLGARD 184-coated perfusion chamber (Automate Scientific, Berkeley, CA) with a bath of AHL-TTX. Perfusion flow was established with a gravity-fed ValveBank II perfusion system (Automate Scientific, Berkeley, CA), and waste was collected with a vacuum pump. Recording was started after the brains had been in AHL-TTX for at least five minutes.

Images were acquired on a Leica Stellaris 8 with confocal imaging equipped with a white-light laser and 405 nm diode. Using LAS X software (Leica, Wetzlar, Germany), we recorded XYZT in Live Imaging Mode. Using a 20X water objective with 0.5 numerical aperture (Leica, Wetzlar, Germany), images were acquired at 512×512 with the pinhole setting at 6.73. CFP was excited with a 440 nm laser and detected using an HyD S2 sensor, while YFP was detected using an HyD S 4 sensor. Laser intensities and detector gains were kept consistent within individual experiments. With both the ITP^+^ LNd and the 5^th^ s-LNv in frame, z-positions were set for both cells. Images of both z-positions were taken every two seconds. Recordings included one minute of baseline recording with AHL-TTX perfusion and three minutes of AHL-TTX with 200 uM dopamine perfusion.

Images were analyzed with custom MATLAB code modified from Adel et al., 2022. CFP and YFP signals were individually extracted using hand-drawn ROIs, and they were normalized to background signal. The ratio CFP/YFP indicates cAMP signal. Then, ΔF/F was calculated, with the average of 10 seconds before the onset of dopamine perfusion acting as a baseline. Python was used to plot the time series with a simple moving average of two data points and with the shaded regions signifying the 68% confidence interval of the mean.

## Notes

### Competing Interest Statement

The authors have declared no competing interest.

