## Supplementary Figures for "Light and dopamine impact two circadian neurons to promote morning wakefulness"

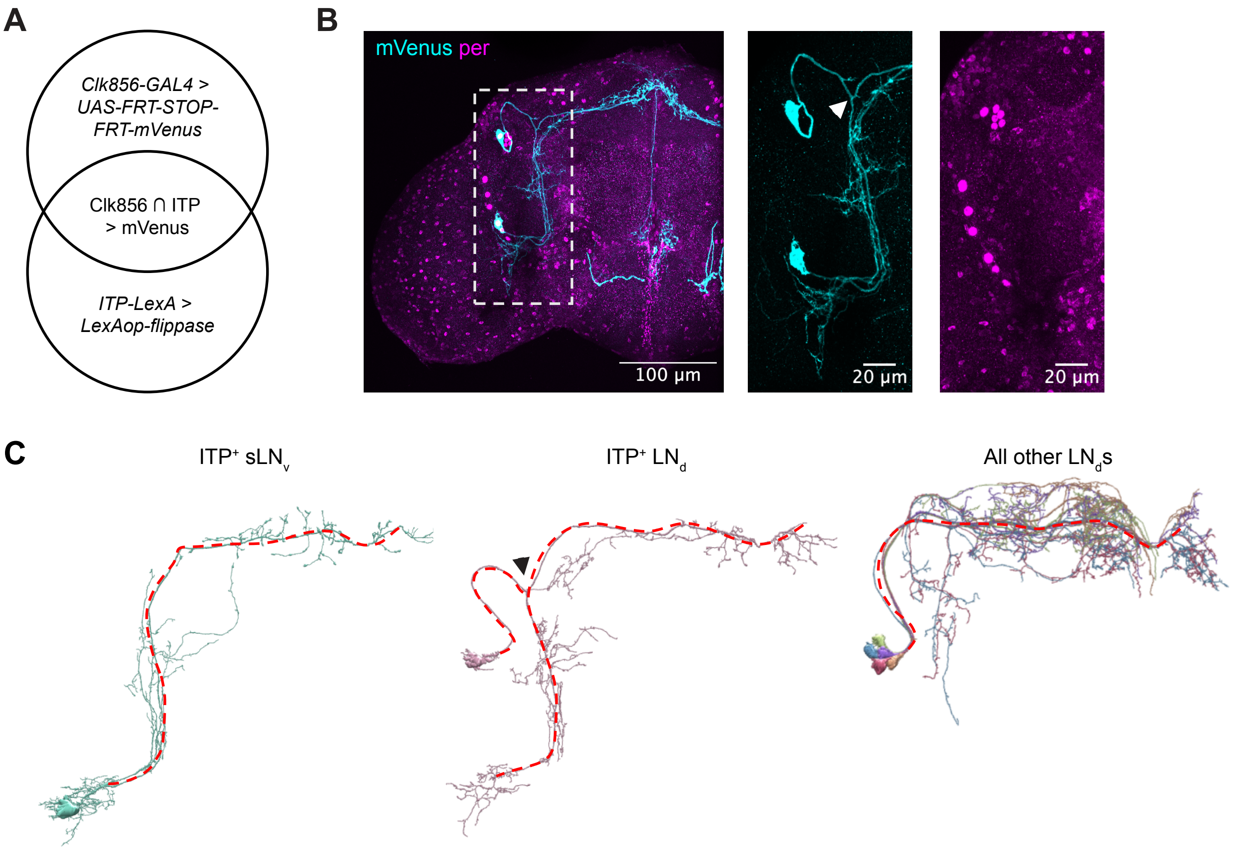


**Figure S1. The ITP+ LN$_d$ is morphologically distinct by a characteristic axonal bifurcation.**

(A) An intersectional strategy to label cells in common to *Clk856-GAL4* and *ITP-LexA* driver lines with mVenus.

(B) Representative images of the intersection of *Clk856-GAL4* and *ITP-LexA* marked by mVenus (cyan) and period (magenta) antibody staining. Center and right images are a close-up of the dashed white box area on the left. White arrowhead marks an axonal bifurcation characteristic of ITP+ LN$_d$s.

(C) EM connectome tracing of the ITP+ LNs and the other LN$_d$s. Axons are marked with red dashed lines. EM reconstructions were visualized using NeuPrint [1].

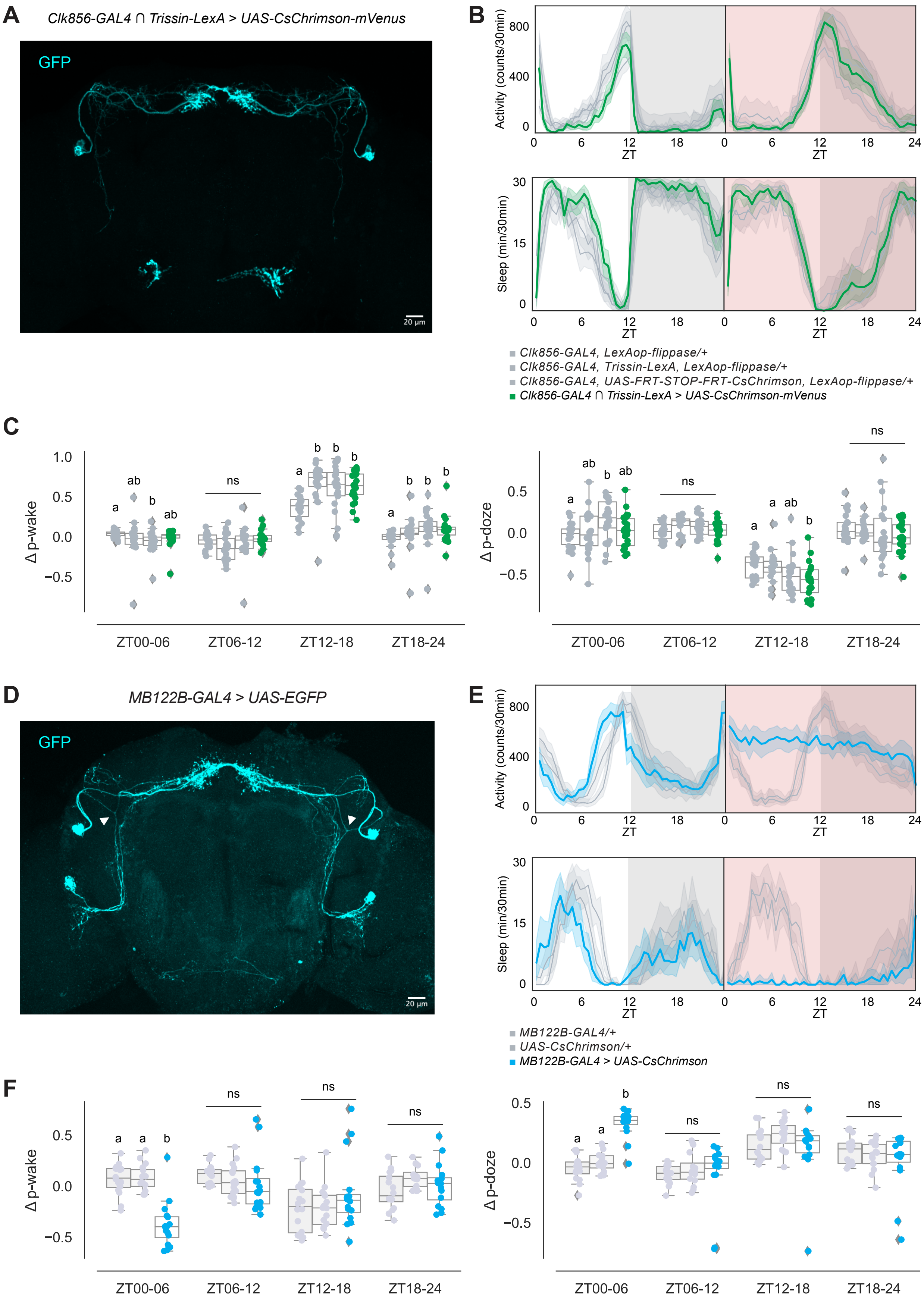
A
Clk856-GAL4 ∩ Trissin-LexA > UAS-CsChrimson-mVenus
GFP
20 µm
B
Activity (counts/30min)
800
400
0
Sleep (min/30min)
30
15
0
0 6 12 18 0 6 12 18 24
ZT
Clk856-GAL4, LexAop-flippase/+
Clk856-GAL4, Trissin-LexA, LexAop-flippase/+
Clk856-GAL4, UAS-FRT-STOP-FRT-CsChrimson, LexAop-flippase/+
Clk856-GAL4 ∩ Trissin-LexA > UAS-CsChrimson-mVenus
C
Δ p-wake
Δ p-doze
1.0
0.5
0.0
−0.5
a ab b ab
ns
a b b b
a b b b
ab b ab
ns
a ab b
ZT00-06
ZT06-12
ZT12-18
ZT18-24
D
MB122B-GAL4 > UAS-EGFP
GFP
20 µm
E
Activity (counts/30min)
Sleep (min/30min)
MB122B-GAL4/+
UAS-CsChrimson/+
MB122B-GAL4 > UAS-CsChrimson
F
Δ p-wake
Δ p-doze
a a b
ns
ZT00-06
ZT06-12
ZT12-18
ZT18-24

**Figure S2. ITP$^+$ LNs and not the Trissin$^+$ LN$_d$s are sufficient for promoting activity.**

(A) Representative whole-brain maximum intensity projection of the intersection of *Clk856-GAL4* and *Trissin-LexA* labeling Trissin$^+$ LN$_d$s with GFP (cyan) antibody staining.

(B) Timeseries plots of activity and sleep of male flies expressing red light-sensitive CsChrimson in Trissin$^+$ LN$_d$s at baseline (left) and with 24-hour red LED optogenetic activation (right) in 12:12 LD. Heterozygous no UAS/LexA, no UAS, and no LexA controls are in gray. Experimental flies expressing all transgenes are colored. Bold lines are means and shaded regions are 95% confidence intervals of the means.

(C) Boxplots quantifying the change in p-wake (left) and p-doze (right) of male flies expressing red light-sensitive CsChrimson in Trissin$^+$ LN$_d$s during the red LED optogenetic activation day from baseline day in six-hour time bins. Genotypes are depicted by the same color scheme as in (B). Letters represent statistically distinct groups as tested by a Kruskal Wallis test, post-hoc Mann Whitney U multiple comparisons method, and a Bonferroni-corrected significance value of $p < 0.008333$. Groups labeled with "ns" are not statistically distinct.

(D) Representative whole-brain maximum intensity projection of *MB122B-GAL4* labeling ITP$^+$ LNs and Trissin$^+$ LN$_d$s with GFP (cyan) antibody staining. Axonal bifurcations of the ITP$^+$ LN$_d$s are marked by white arrowheads.

(E) Timeseries plots of activity and sleep of female flies expressing red light-sensitive CsChrimson in ITP$^+$ LNs and Trissin$^+$ LN$_d$s at baseline (left) and with 24-hour red

LED optogenetic activation (right) in 12:12 LD. Heterozygous *MB122B-GAL4* and *UAS-CsChrimson* controls are in gray. Experimental flies expressing both transgenes are colored. Bold lines are means and shaded regions are 95% confidence intervals of the means.

(F) Boxplots quantifying the change in p-wake (left) and p-doze (right) sleep of female flies expressing red light-sensitive CsChrimson in $ITP^+$ LNs and $Trissin^+$ $LN_d$s during the red LED optogenetic activation day from baseline day in six-hour time bins. Genotypes are depicted by the same color scheme as in (E). Letters represent statistically distinct groups as tested by a Kruskal Wallis test, post-hoc Mann Whitney U multiple comparisons method, and a Bonferroni-corrected significance value of $p < 0.01667$. Groups labeled with "ns" are not statistically distinct.

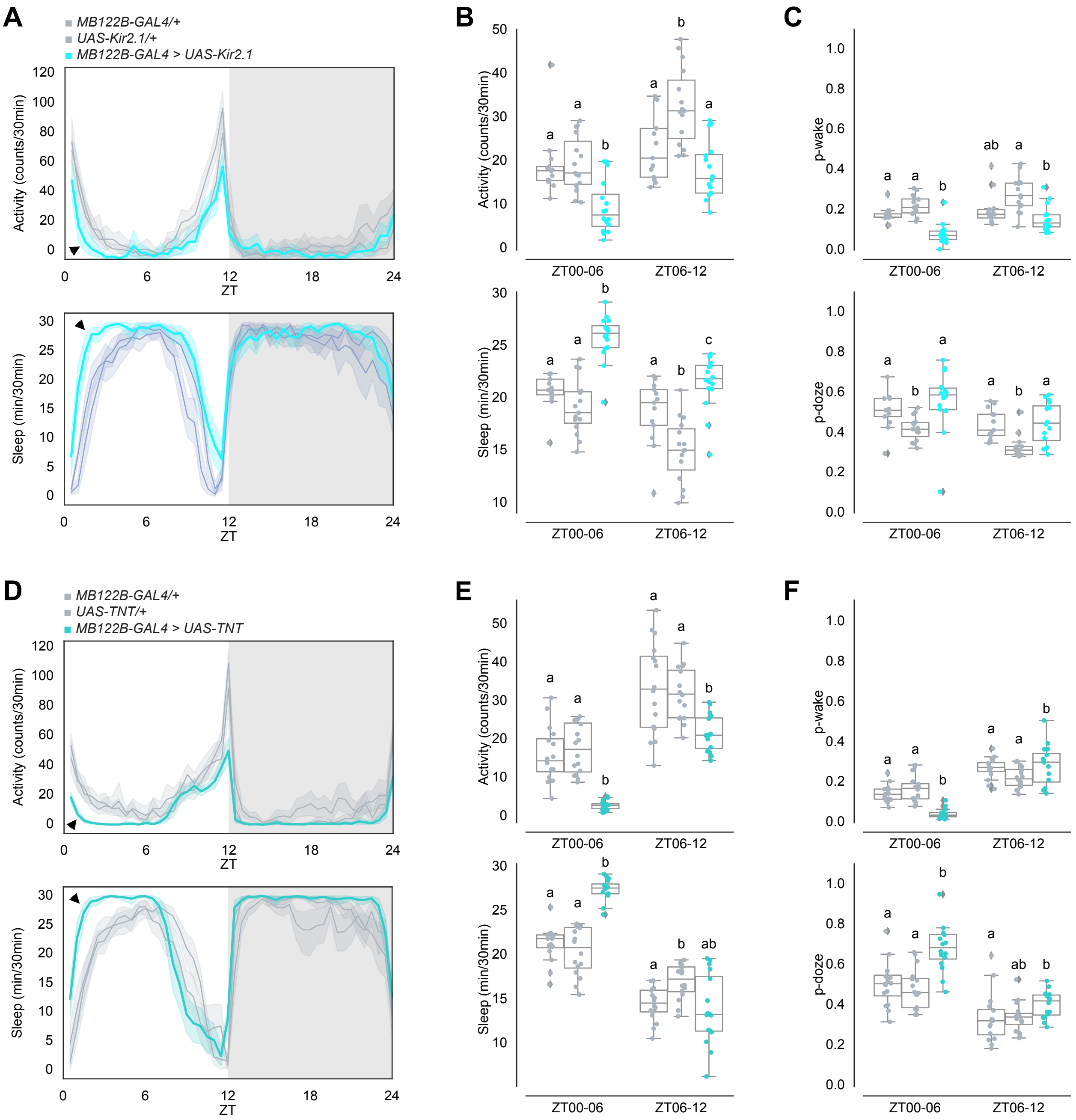


**Figure S3. Inhibiting $ITP^+$ LNs and $Trissin^+$ $LN_d$s decreases early morning wakefulness.**

(A) Timeseries plots of activity (top) and sleep (bottom) of male flies expressing inhibitory Kir2.1 potassium channels in $ITP^+$ LNs and $Trissin^+$ $LN_d$s in 12:12 LD.

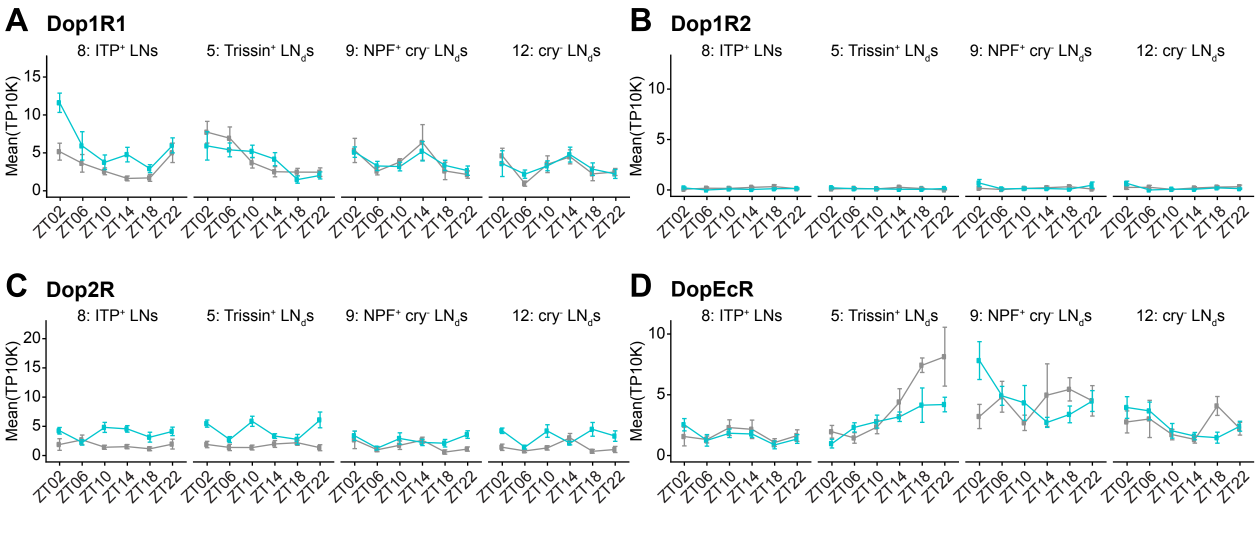


**Figure S4. *Dop1R1* mRNA cycles in 12:12 LD in ITP⁺ LNs but not in the other $LN_d$s.**

(A) Timeseries plots of *Dop1R1* mRNA expression levels around the clock in distinct evening cell subtypes from previously published single-cell RNA sequencing data [2]. Blue lines are mRNA levels in 12:12 LD and gray lines are mRNA levels in DD. Lines are mean values in number of transcripts per ten thousand transcripts and bars represent the standard error of the mean.

(B) Timeseries plots of *Dop1R2* mRNA expression levels around the clock in distinct evening cell subtypes from previously published single-cell RNA sequencing data [2]. Blue lines are mRNA levels in 12:12 LD and gray lines are mRNA levels in DD. Lines are mean values in number of transcripts per ten thousand transcripts and bars represent the standard error of the mean.

(C) Timeseries plots of *Dop2R* mRNA expression levels around the clock in distinct evening cell subtypes from previously published single-cell RNA sequencing data [2]. Blue lines are mRNA levels in 12:12 LD and gray lines are mRNA levels in

DD. Lines are mean values in number of transcripts per ten thousand transcripts and bars represent the standard error of the mean.

(D) Timeseries plots of *DopEcR* mRNA expression levels around the clock in distinct evening cell subtypes from previously published single-cell RNA sequencing data [2]. Blue lines are mRNA levels in 12:12 LD and gray lines are mRNA levels in DD. Lines are mean values in number of transcripts per ten thousand transcripts and bars represent the standard error of the mean.

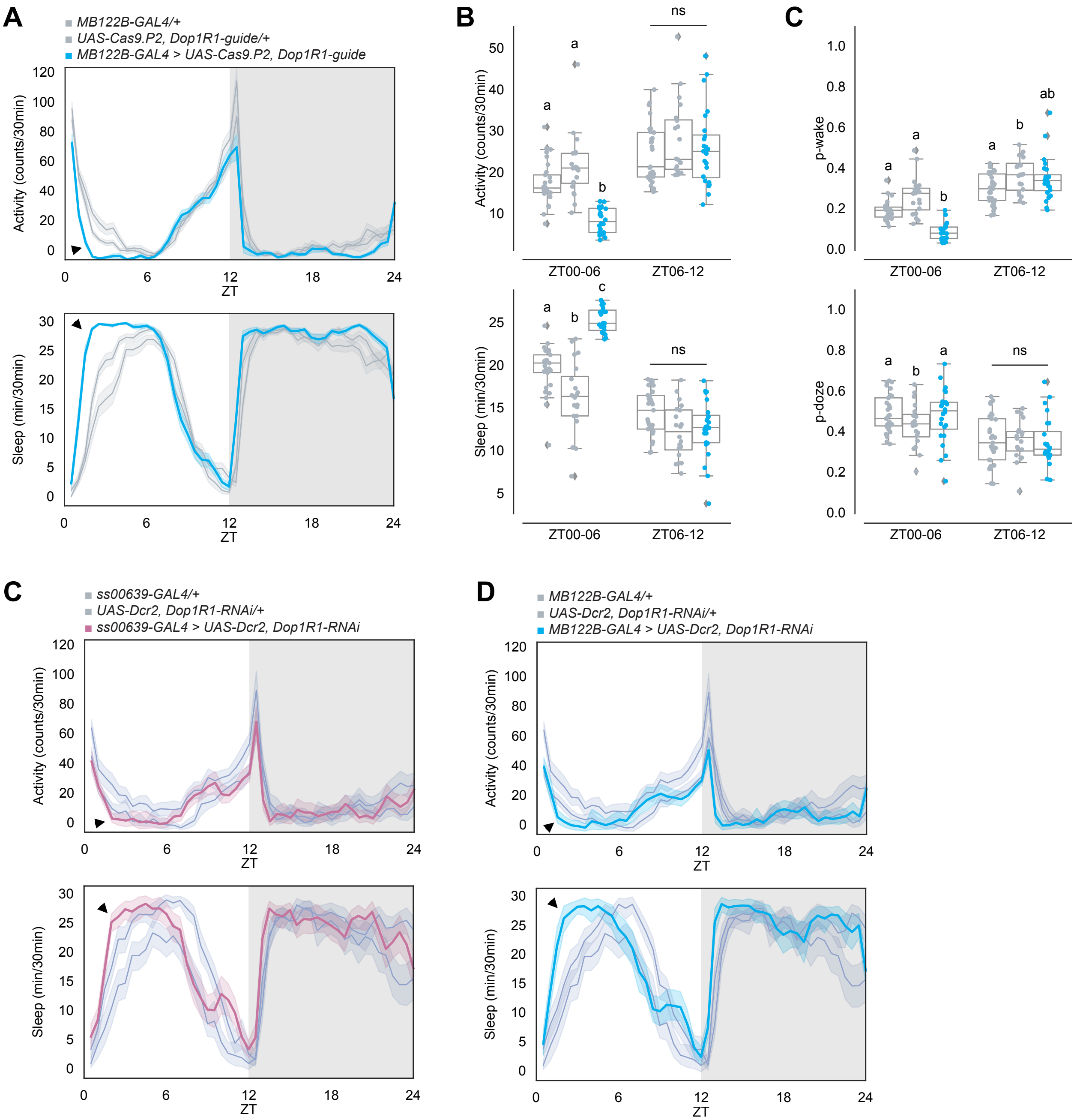


**Figure S5. RNAi Knockdown of Dop1R1 in ITP⁺ LNs lead to a small deficit in early morning wakefulness.**

(A) Timeseries plots of activity (top) and sleep (bottom) of male flies with *Dop1R1* knock-out in ITP⁺ LNs and Trissin⁺ LN$_d$s in 12:12 LD. Heterozygous *MB122B-*

*GAL4* and *UAS-Cas9.P2, Dop1R1-guide* controls are in gray. Experimental flies expressing all transgenes are colored. Bold lines are means and shaded regions are 95% confidence intervals of the means. Black arrowheads point to the major difference between control and experimental flies between ZT00 and ZT06.

(B) Boxplots quantifying activity (top) and sleep (bottom) of male flies with *Dop1R1* knock-out in ITP$^+$ LNs and Trissin$^+$ LN$_d$s during ZT00-06 and ZT06-12 daytime bins. Genotypes are depicted by the same color scheme as in (A). Letters represent statistically distinct groups as tested by a Kruskal Wallis test, post-hoc Mann Whitney U multiple comparisons method, and a Bonferroni-corrected significance value of $p < 0.01667$. Groups labeled with “ns” are not statistically distinct.

(C) Boxplots quantifying p-wake (top) and p-doze (bottom) of male flies with *Dop1R1* knock-out in ITP$^+$ LNs and Trissin$^+$ LN$_d$s during ZT00-06 and ZT06-12 daytime bins. Genotypes are depicted by the same color scheme as in (A). Letters represent statistically distinct groups as tested by a Kruskal Wallis test, post-hoc Mann Whitney U multiple comparisons method, and a Bonferroni-corrected significance value of $p < 0.01667$. Groups labeled with “ns” are not statistically distinct.

(D) Timeseries plots of activity (top) and sleep (bottom) of female flies with *Dop1R1* knock-down in ITP$^+$ LNs in 12:12 LD. Heterozygous *ss00639-GAL4* and *UAS-Dcr2, Dop1R1-RNAi* controls are in gray. Experimental flies expressing all transgenes are colored. Bold lines are means and shaded regions are 95%

confidence intervals of the means. Black arrowheads point to the major difference between control and experimental flies between ZT00 and ZT06.

(E) Timeseries plots of activity (top) and sleep (bottom) of female flies with *Dop1R1* knock-down in ITP$^+$ LNs and Trissin$^+$ LN$_d$s in 12:12 LD. Heterozygous *MB122B-GAL4* and *UAS-Dcr2, Dop1R1-RNAi* controls are in gray. Experimental flies expressing all transgenes are colored. Bold lines are means and shaded regions are 95% confidence intervals of the means. Black arrowheads point to the major difference between control and experimental flies between ZT00 and ZT06.

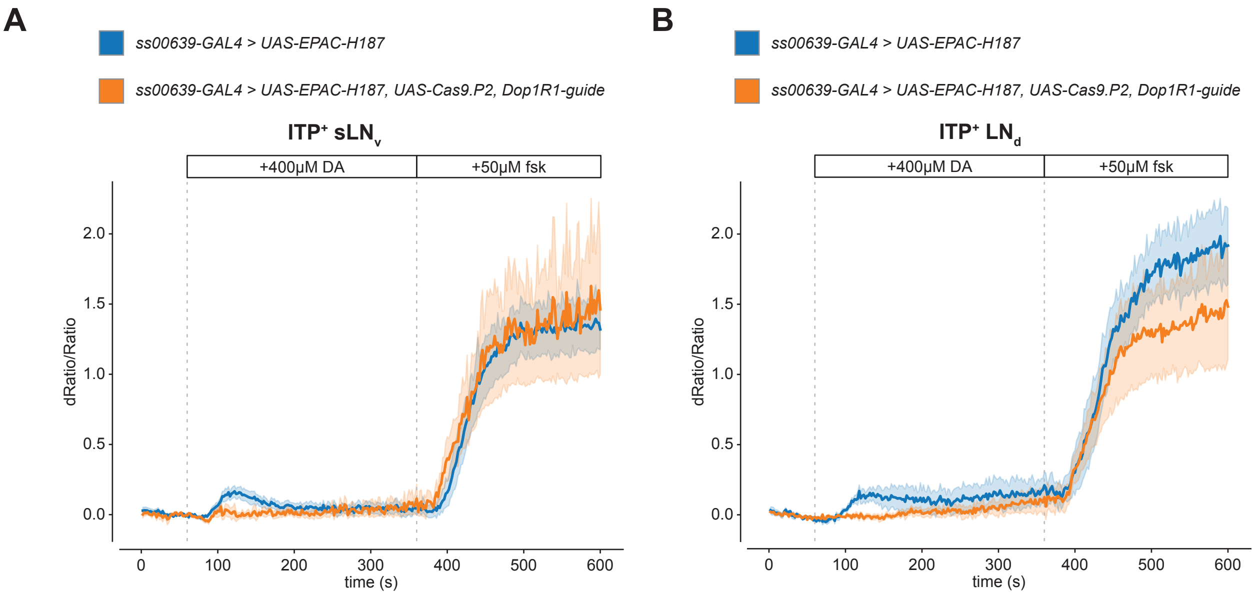


**Figure S6. cAMP responses to dopamine are dependent on Dop1R1 expression in ITP⁺ LNs.**

(A) Timeseries plot of cAMP levels at baseline and in response to 400μM dopamine and 50μM forskolin perfusion as measured by *UAS-EPAC-H187* in the ITP⁺ $sLN_v$ with (orange) and without (blue) Dop1R1 knock-out using *UAS-Cas9.P2, Dop1R1-guide*. Bold lines are means shown with a simple moving average of two data points and shaded regions are 68% confidence intervals of the means.

(B) Timeseries plot of cAMP levels at baseline and in response to 400μM dopamine and 50μM forskolin perfusion as measured by *UAS-EPAC-H187* in the ITP⁺ $LN_d$ with (orange) and without (blue) Dop1R1 knock-out using *UAS-Cas9.P2, Dop1R1-guide*. Bold lines are means shown with a simple moving average of two data points and shaded regions are 68% confidence intervals of the means.
